# Dynamic prospect theory - two core economic decision theories coexist in the gambling behavior of monkeys

**DOI:** 10.1101/2021.04.04.438415

**Authors:** Agnieszka Tymula, Yuri Imaizumi, Takashi Kawai, Jun Kunimatsu, Masayuki Matsumoto, Hiroshi Yamada

## Abstract

Research in behavioral economics and reinforcement learning has given rise to two influential theories describing human economic choice under uncertainty. The first, prospect theory, assumes that decision-makers use *static* mathematical functions, utility and probability weighting, to calculate the values of alternatives. The second, reinforcement learning theory, posits that *dynamic* mathematical functions update the values of alternatives based on experience through reward prediction error (RPE). To date, these theories have been examined in isolation without reference to one another. Therefore, it remains unclear whether RPE affects a decision-maker’s utility and/or probability weighting functions, or whether these functions are indeed static as in prospect theory. Here, we propose a dynamic prospect theory model that combines prospect theory and RPE, and test this combined model using choice data on gambling behavior of captive macaques. We found that under standard prospect theory, monkeys, like humans, had a concave utility function. Unlike humans, monkeys exhibited a concave, rather than inverse-S shaped, probability weighting function. Our dynamic prospect theory model revealed that probability distortions, not the utility of rewards, solely and systematically varied with RPE: after a positive RPE, the estimated probability weighting functions became more concave, suggesting more optimistic belief about receiving rewards and over-weighted subjective probabilities at all probability levels. Thus, the probability perceptions in laboratory monkeys are not static even after extensive training, and are governed by a dynamic function well captured by the algorithmic feature of reinforcement learning. This novel evidence supports combining these two major theories to capture choice behavior under uncertainty.

**Significance statement:** We propose and test a new decision theory under uncertainty by combining pre-existing two influential theories in the neuroeconomics: prospect theory from economics and prediction error theory from reinforcement learning. Collecting a large dataset (over 60,000 gambling decisions) from laboratory monkeys enables us to test the hybrid model of these two core decision theories reliably. Our results showed over-weighted subjective probabilities at all probability levels after lucky win, indicating that positive prediction error systematically bias decision-makers more optimistically about receiving rewards. This trial-by-trial prediction-error dynamics in probability perception provides outperformed performance of the model compared to the standard static prospect theory. Thus, both static and dynamic elements coexist in monkey’s risky decision-making, an evidence contradicting the assumption of prospect theory.

## Introduction

The multidisciplinary field of neuroeconomics (1-3) has made significant progress towards understanding how the brain makes economic choices. By combining both theoretical and empirical tools from neuroscience, psychology, and economics, the long-term goal of neuroeconomics is directed to construct a biologically viable, unified framework explaining economic choice. Two models — prospect theory (4) and reinforcement learning theory (5) — have been particularly popular and tested at both the behavioral and neural levels, although these models have predominantly been studied in isolation from one another. Prospect theory captures static features of risky behaviors through fixed preference parameters. Contrastingly, reinforcement learning focuses on learning the value of rewards, which is the dynamic aspect of decision-making. These two separate theories have sometimes described similar behavioral and neural observations related to risky decision-making, but we do not yet know how these two theories come together to explain the decision-making. In particular, we do not know whether reward prediction error, an algorithmic feature of reinforcement learning, affects the utility of rewards or perception of the probability with which rewards are received. While the consensus view is that both theories describe risky decision-making well under typical experimental conditions, a joint exploration of these models has not been conducted, even at the behavioral level (6). This is because joint examination of these two theories requires reliable estimate of many parameters from the two models, but obtaining large datasets sufficient for the estimation is difficult in human subjects. In this paper, we overcome this issue by collecting behavioral data from two monkeys to build a sufficiently large dataset, which allows us to explore a model that combines both theories.

### Prospect theory

Economic theory has developed well-specified, rigorous models of human choice under uncertainty (7-10). There is a broad consensus in economics that humans capture the desirability of a reward through a utility function and that, if rewards are probabilistic, they calculate expected utilities by multiplying utilities of rewards by the probabilities with which they are received (11). Prospect theory (5) extends this framework by allowing for the subjective perception of probabilities through a static, inverse-S shaped probability weighting function, in which small probabilities are overestimated and large probabilities are underestimated (4, 12). Utility functions have been estimated in hundreds of behavioral studies in humans and have typically been found to be concave in the domain of gains. Probability weighting functions estimated on the aggregate level in humans are typically found to be inverse-S shaped (11, 12). However, weighting functions estimated at the individual subject level differ between individuals and often take shapes far from the canonical inverse-S (13-19). While this individual heterogeneity may arise from many reasons, one possible factor would be a fluctuation of probabilistic events experienced individually. This possibility poses a challenge for theory, which is yet to explain the change in probability perception at the individual level.

The expected utility and prospect theory models of decision-making have been used to study animal behavior in different disciplines (3, 20). Due to the nature of tasks that animals can undertake, these studies have been limited to the domain of gains. In this domain, most humans exhibit a concave utility function, consistent with risk-averse behavior. The utility functions of many species — bees, birds, rodents, and non-human primates — have been estimated, but with inconsistent and sometimes controversial results (20-24). Studies in laboratory monkeys, a standard neurobiological model for human economic choice, have similarly yielded conflicting results. Some studies found that, unlike humans, rhesus monkeys exhibit a convex utility function under typical experimental conditions (24-28). However, under other experimental conditions, these same animals behave as if they have a concave or linear utility (25, 28). Other studies have found concave or S-shaped utility functions in monkeys (29-32) similar to humans, consistent with the notion that monkeys are an appropriate model for the study of human choice behavior. The prospect theory have begun to ask whether captive macaques also distort probabilities in the same way as humans are believed to and have arrived at inconsistent conclusions (26, 27, 32-34).

### Reinforcement learning theory

Significant progress has also been made towards understanding how animals learn the value of available rewards for choice behavior through experience (5). In reinforcement learning theory, unlike under the economic approach, the individual may not know the precise value or utility of all the alternatives under consideration. The reinforcement learning model specifies how the individual dynamically updates the value of each item or option to make choices by comparing the value of obtained reward with its predicted value via reward prediction error (RPE) (35, 36). This mathematical algorithm, RPE, captures learning values and has been extensively examined in choice behavior and its neural basis of humans, monkeys, and rodents under uncertainty (37-43). However, whether the mechanism for reinforcement learning involves updating the utility of reward or its probability weighting has not been established.

### Combined approach in the present study

We combine the approaches of static prospect theory and the dynamic reinforcement learning theory into a hybrid model, which includes the critical features of both models — utility function, probability weighting function, and RPE. To test the empirical validity of this combined model, which we call dynamic prospect theory, we designed an experiment in which monkeys choose gamble options, in which the probabilities and magnitudes of rewards were completely orthogonal. We estimated the parameters of the hybrid model reliably with a large number of choice trials collected in each individual monkeys, which is impossible in human subjects. Our estimation revealed that the probability weighting function, but not the curvature of the utility function, varies with the RPE, indicating that probability weighting is dynamically adjusted decision-by-decision, contradicting the assumption in static prospect theory.

## Results

To determine the monkeys’ internal valuation processes, we trained them to perform a gambling task (Figure 1A) (44), similar to previous experiments performed in human subjects in economics (45) and reinforcement learning (46). The task involved choosing between two options, each offering an amount of liquid reward with a probability. The monkeys fixated on a central gray target. Then, two options were presented visually as pie-charts displayed on the left and right sides of the screen. The number of green pie segments indicated the magnitude of liquid reward in 0.1 mL increments (0.1-1.0 mL) and the number of blue pie segments indicated the probability of receiving the reward in 0.1 increments (0.1-1 where 1 indicates a 100% chance). After the pie-chart disappeared, the gray target in the center reappeared for 0.5 seconds. Thereafter, the monkeys chose between the left and right targets by fixating on one of the sides. Following the choice, the monkeys received or did not receive the amount of liquid reward associated with their chosen option according to its corresponding probability, otherwise no-reward. In each choice trial, two out of the 100 possible combinations of the reward probability and magnitude (Figure 1B) were randomly allocated to the left- and right-side target options. We aggregated all data collected after each monkey learned to associate the probability and magnitude with the pie-chart stimuli. There were 44,883 decisions by monkey SUN (obtained in 884 blocks over 242 days) and 19,292 decisions by monkey FU (obtained in 571 blocks over 127 days). Well-trained monkeys, like humans, showed behavior consistent with utility maximization, selecting on average options with the higher expected value (Figure 1C).

**Fig. 1.**
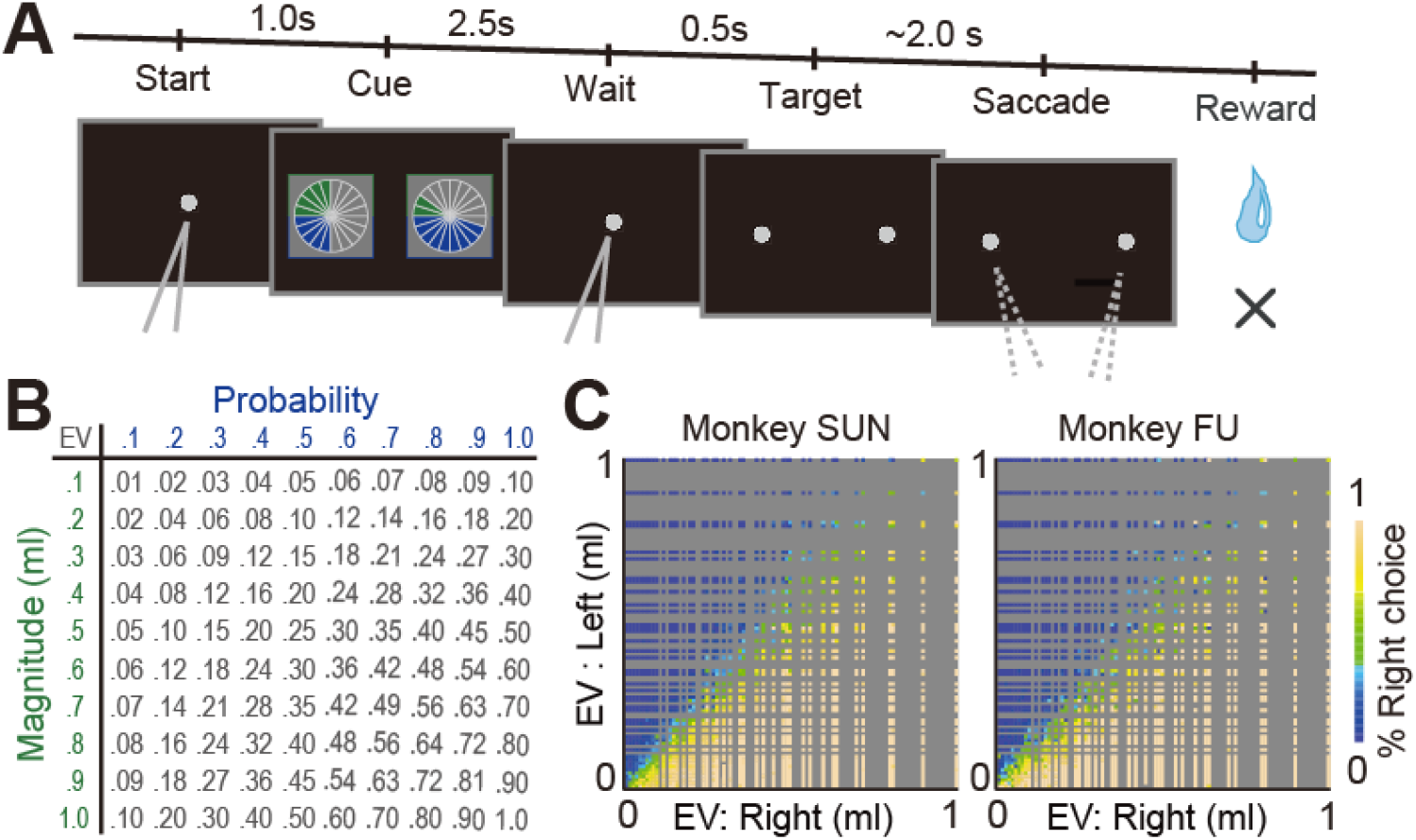
Cued lottery task and monkeys’ choice behavior. (A) A sequence of events in choice trials. Two pie charts representing the available options were presented to the monkeys on the left and right sides of the screen. Monkeys chose either of the targets by fixating on the side where it appeared. (B) Payoff matrix – each magnitude was fully crossed with each probability, resulting in a pool of 100 lotteries from which two were randomly allocated to the left- and right-side target options on each trial. Expected values (EVs) are presented in mL. (C) Frequency with which the target on the right side was selected for the expected values of the left and right target options.

### Estimation of static prospect theory parameters

We first estimated each monkey’s utility and probability weighting functions using standard parametrizations in the literature. For the utility function, we used the power utility function *U(m) = m*^*α*^, where *m* indicates the magnitude of the reward, *α*>1, indicates convex utility (risk-seeking), *α*<1 indicates concave utility (risk aversion), and *α*=1 indicates linear utility (risk neutrality). Overall, we estimated the following five sequentially developed models of the utility of receiving reward magnitude *m* with probability *p, V(p,m)*:

1. EV: expected value *V(p,m) = p* × *m*
2. EU: expected utility *V(p,m)* = *p* × *m*^*α*^
3. TK92: prospect theory with *w(p)* as in *(4) V(p,m)* = *p*^*γ*^/ (*p*^*γ*^+(1-*p*)^*γ*^)^1/*γ*^ × *m*^*α*^
4. Prelec: prospect theory with *w(p)* as in (47) *V(p,m)* = *exp(-δ (-log p)* ^*γ*^) × *m*^*α*^
5. GE: prospect theory with *w(p)* as in (48) *V(p,m)* = *δp*^*γ*^ / (*δp*^*γ*^+(1-*p*)^*γ*^) × *m*^*α*^

Where *α, γ* and *δ* are free parameters and *p* and *m* are the probability and magnitude cued by the lottery, respectively. To determine which model best describes observed monkey’s behavior, we compared the Baisian Information Criterion (BIC) term for each model, the goodness of model fit with a penalty (see Methods for more details). The smallest BIC values among the five models showed that Prelec and GE models outperformed EV, EU, and TK92 (Fig. 2A and Table S1)

**Fig. 2.**
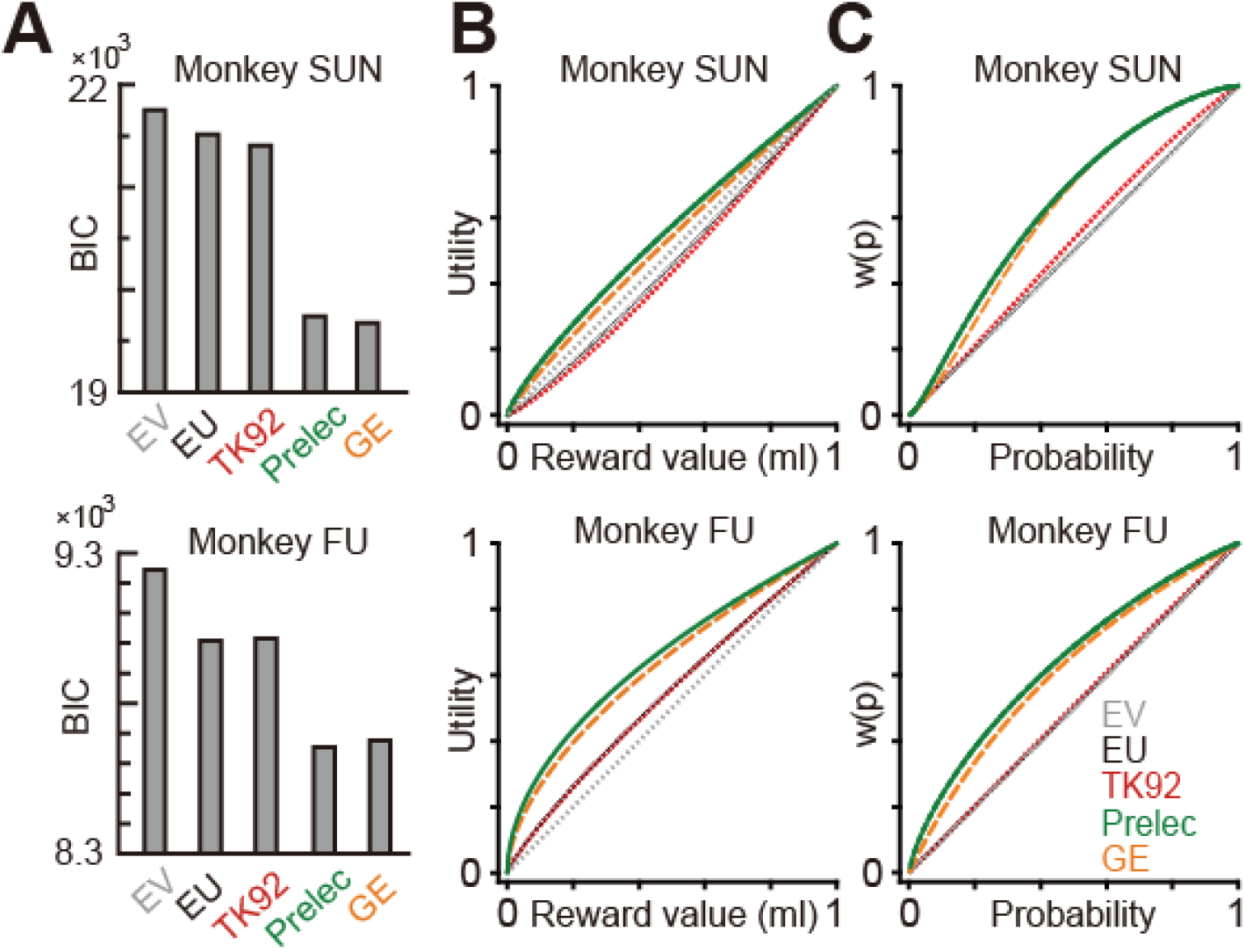
Estimated utility and probability weighting function in monkeys (A) BIC values of the standard economic models: EV, EU, TK92, Prelec, and GE. See methods for details. (B) Drawing of the estimated utility functions in different models. (C) Drawing of the estimated probability weighting functions in different models.

The curvature of the utility function was predominantly concave (Fig. 2B, see green, Prelec and orange, GE). Notably, for both monkeys, the two-parameter probability weighting functions were concave, instead of the inverse-S shape traditionally assumed for in humans (Fig. 2C, see green and orange). We further established the robustness of this observation using simple logistic regression analysis. Both monkeys chose lotteries with probabilities below 0.3 and above 0.7 less often, relative to lotteries with middle range probabilities (Table S2). Such a choice pattern is consistent only with a concave probability weighting function. If the monkeys had an inverse-S probability weighting function, they would have chosen lotteries with low probabilities more often and lotteries with high probabilities less often than lotteries with middle range probabilities.

The estimates from the Prelec and GE models are remarkably different from the other models. In particular, the EU and TK92 models yield more convex (in monkey SUN) and less concave (in monkey FU) utility functions (Fig. 2B, see black and red). Note that probability distortions are absent by the assumption in EU (Fig. 2C, black). In TK92, the probability weighting function can only capture inverse-S shaped or linear functions, and thus, the estimated probability distortions are either absent (monkey FU) or very slight (monkey SUN) (Fig. 2C, red, Table S1, Gamma parameter in TK92). Since EU and TK92 models cannot capture increased risk-taking that stems from concave probability weighting, the EU and TK92 models require more convex / less concave utility functions. Nevertheless, both the EU and TK92 models yield similar goodness of fit, better than those with the EV model (Fig. 2A).

Overall, our orthogonal data matrix with most flexible model of probability distortion leads to the conclusion that monkeys distort probability different from that usually assumed for human decision-makers. Further, when subjective distortions in probability are accounted for, monkeys’ estimated utility functions are concave.

### Reinforcement learning model illustration

We illustrate how the typical reinforcement learning approach would model monkey’s valuation in our experimental condition (see methods). In each trial *t* for each monkey, they received a lottery outcome according to their chosen lottery, described by a combination of probability and magnitude. After receiving the reward or not, the value function is updated through a learning algorithm *V(p,m)* _*t+1*_ *= V(p,m)*_*t*_ *+ AΔ*_*t*_, where *Δ*_*t*_ *= r*_*t*_ *- V(p,m)*_*t*_ is the reward prediction error (the difference between the received reward, *r*_*t*_, and the predicted value of the lottery, *V(m,p)*_*t*_). *A* is the learning rate at which the value function is updated ^***1***^. We simulated this reinforcement learning model using different learning rates for each of the 100 lotteries for 10,000 trials. After 10,000 trials, the algorithm produced the lottery valuations, *V(m,p)*_*t=10,000*_, that were reasonable given the learning rates (Fig. S1A). At lower learning rates, which would usually be observed in stable environments like this experiment, these valuation functions arrived at predicted lottery valuations that are very close to the expected value., i.e., probability times magnitude. There were slight deviations in the predictions from the expected value (diagonal line) due to the RPE, which yielded trial-by-trial dynamics in *V(p,m)*_*t*_ even after substantial learning. The extent of these fluctuations was causally related to the learning rate (Fig. S1A) and these fluctuations naturally existed with both positive (Fig. S1B, reward) and negative (Fig. S1C, no-reward) RPE.

This simple simulation exercise demonstrates how the typical reinforcement learning model captures the gambling behavior of the monkeys in our experiment, although the reinforcement learning model does not reveal whether the utility curvature and/or subjective probabilities are influenced by RPE. Thus, under the reinforcement learning approach alone, it remains unclear whether, after receiving a reward that is larger than predicted, the reward itself becomes more valuable or whether an individual’s belief about the probability of receiving the reward increases.

### Dynamic prospect theory model

To answer the above question, we propose a dynamic prospect model which combines the elements of both reinforcement learning theory and prospect theory into a single framework. In doing so, we made four assumptions. First, we used Prelec as our baseline model, as it best fit the data. Second, based on previous studies (49, 50), we allowed for positive and negative reward prediction errors to differently affect parameters in our combined model. Third, we allowed the reward prediction error to affect the parameters of utility and probability weighting functions, so that we could estimate where significant effects occur. Fourth, given our model simulation (Fig S1A), that shows that after substantial learning, monkeys should have arrived at relatively stable predicted valuations of lotteries well captured by expected value on average, we made a simplifying assumption that *Δ*_*t*_ *= r*_*t*_ *- EV(p,m)*_*t*_. This implies that in our setting, a positive reward prediction error always occurs after the reward was received and it is bigger the larger the reward amount was and the lower was the probability of receiving it. A negative reward prediction error always occurs in trials where the reward is not received, and it is smaller (more negative) the larger the reward amount was and the larger was the probability with which the reward could be received. Our dynamic prospect theory model has dynamic, rather than static, parameters:

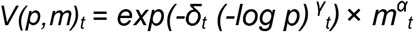

We assumed that the dynamic parameters of this model could be affected, in principle, by three variables: a positive reward prediction error after receiving a reward (*Δ*positive_*t-1*_), a negative reward prediction error after a trial when a reward was not received (*Δ*negative_*t-1*_), and an indicator variable denoting whether the monkey received the reward or not in the past trial (fb_*t-1*_). We captured these effects by rewriting the model parameters as follows:

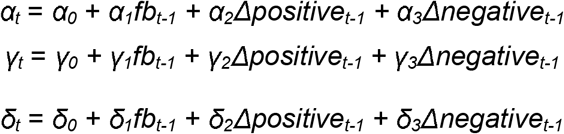

Our hybrid model can identify both the static and the dynamic nature of the internal valuation process. If only the constant parameters, *α*_*0*_, *γ*_*0*_, and *δ*_*0*_, are significant, the model collapses into the traditional static prospect theory model. If the parameters on the past trial variables, *α*_*1*_*-α*_*3*_, *γ*_*1*_*-γ*_*3*_, and *δ*_*1*_*-δ*_*3*_, are significant, this points towards a dynamic model in which valuation adjusts in a way consistent with the ideas behind the reinforcement learning model.

### Empirical test of the dynamic prospect theory model

Our dynamic prospect theory model fits data better than the best-fitting static model (Fig. 3A). We found a clear dependence of the probability weighting function parameter delta (which controls curvature when gamma is close to one) on the positive reward prediction error in both monkeys (Table S3). The larger the positive reward prediction error, the more concave the probability weighting function (Fig. 3B – compare solid green and gray curves). This suggests a more optimistic perception of reward probability after an unexpected positive outcome, such as good fortune or jackpot. Remarkably, the utility function itself was not affected by the reward prediction error (See Alpha parameters in Table S3).

**Fig. 3.**
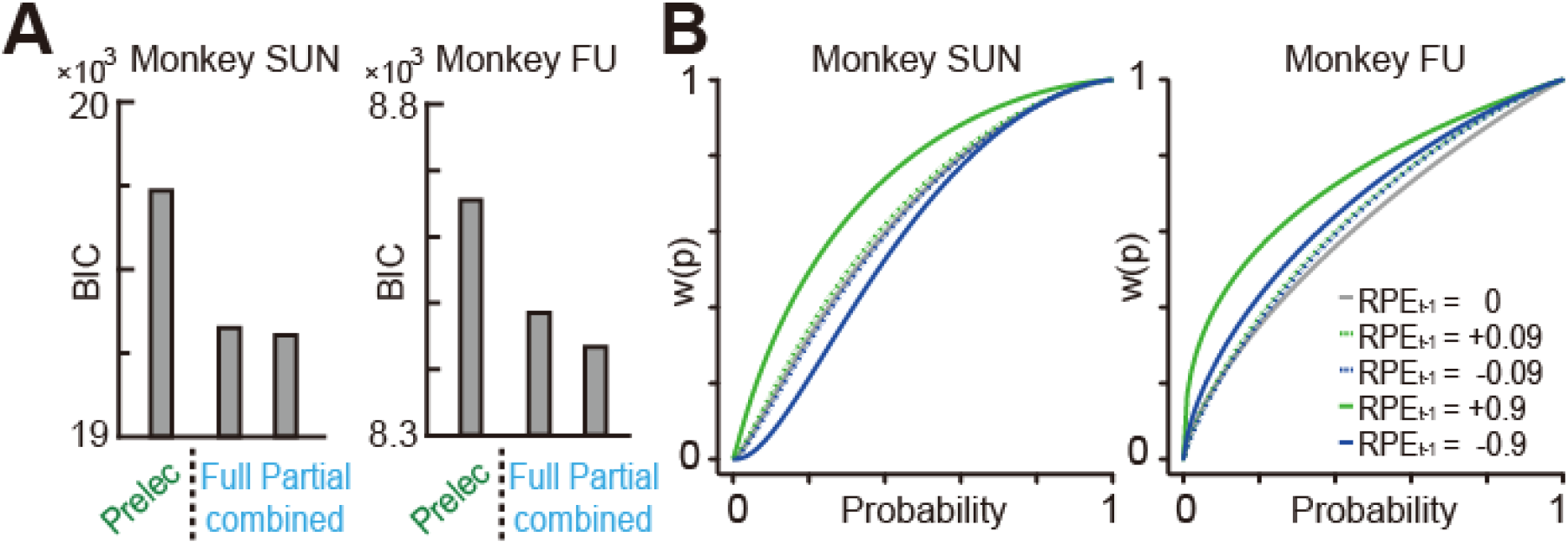
Probability distortion systematically affected by the reward prediction error (A) BIC values estimated in Prelec without reinforcement learning (RL) parameters (Prelec), Prelec with all RL parameters (Full), and Prelec with RL parameters from model selection procedure (Partial). (B) Drawing of the effect of reward prediction error on probability weighting functions in the full combined model, which has been illustrated for trials with the following reward prediction errors: +0.09 in the trials following reward with 0.1 probability and 0.1 ml magnitude; −0.09 in the trials following no-reward with 0.9 probability and 0.1 ml magnitude; +0.9 in the trials following reward with 0.1 probability and 1.0 ml magnitude; −0.9 in the trials following no-reward with 0.9 probability and 1.0 ml magnitude; 0 in the trials following rewards with 100% probability trials.

Because the large number of free parameters in the model may have reduced the accuracy of parameter estimation, we adopted a model selection approach to explore we arrived at the combinations of parameters with the lowest BIC. This approach supports the robustness of our findings. In our best-fitting model (henceforth “partial”, Table S4), we replicated the same effect of RPE on delta and found no effects on utility curvature. Additionally, through the delta parameter, monkey SUN exhibits more pessimistic beliefs about the likelihood of receiving a reward after experiencing larger negative RPE (negative coefficient value for *Δnegative*_*t-1*_, see also Fig. 3B, left, blue). In contrast, this effect is reversed for monkey FU (positive coefficient value for *Δnegative*_*t-1*_, see also Fig. 3B, right, blue). In this best fitting model, we also observe that merely receiving a reward increased gamma, the parameter that controls probability weighting function curvature in terms of the inflection and elevation.

Interpreting these results together, we found that it was the perception of probability, rather than the utility function, that was affected by the reward prediction error signal. This indicates that even after substantial training, the monkeys dynamically updated the subjective probabilities with which they expected to receive rewards, rather than the utility of rewards. These findings may provide an explanation for the “jackpot effect” in gambling behavior.

## Discussion

### Reinforcement learning and probability weighting

We adopted a novel framework to test the integration of the reinforcement learning model into static parameters of utility and probability weighting functions. A priori, we hypothesized that the reinforcement learning algorithm drives a decision-maker’s valuation of gambles either or both through changes in the utility function (subjective value of the reward) and through changes in the subjective probability perception of receiving the reward. Our empirical testing indicate that the probability weighting function dynamically adjusts based on outcome experiences on a trial-by-trial basis, while the utility function remains unaffected. Despite extensive training with the lottery task, the monkeys remained alert to reward probability distributions and updated them through experience. Our most robust finding is that the larger the positive reward prediction error was, the more optimistic the monkeys became about the probability of receiving a reward going forward. Since RPE signal in the brain develops optimal behaviors through the release of dopamine to target brain regions for learning (37, 42, 43, 51), this dynamic feature in the probability perception may or may not be beneficial in our stable experiment, but it may enhance animals to maximize utility in a broader decision-making context when the environment is less stable. It is thought-provoking that we found that the RPE affected only subjective beliefs about probability but not the utility of rewards. Previous empirical studies have found that probability weighting functions systematically vary by age, gender (14), mood (17), and that variations in subjective probabilities could potentially be related to neural coding constraints (52, 53). Our finding adds to this literature in terms of the RPE effect on subjective probabilities, but idiosyncratic in terms of the decision-by-decision dynamics. It is still unclear whether changes in subjective probabilities is affected by an individual’s level of satiety or financial wealth, though existing literature established the effect of satiety on utility functions (21, 22, 30, 54, 55). An interesting extension of this work would be to investigate whether human participants exhibit the same dynamic effects in probability perception when probabilities are precisely communicated with given numerical values.

### Probability weighting of monkeys

A handful of recent studies of captive macaques have begun to investigate static distortions in the perception of probabilities in monkeys, with inconsistent results across studies (26-28, 32-34). The probability weighting function has been found to be inverse-S shaped (27, 33), S-shaped (26, 34), and concave (31, 33) when estimated from economic choice tasks, and either concave or convex when estimated from non-choice tasks(56). Although we consistently found that the probability weighting functions of our two well-trained monkeys were concave, most studies conducted in humans have found inverse-S shaped probability weighting functions on the aggregate level, with a large amount of heterogeneity at the individual level (14, 15, 17-19, 57, 58), with the effects of mood (17), age, and gender (14). However, these studies have not provided a mechanistic explanation about why these heterogeneities exist. Overall, many reasons could explain these inconsistencies in findings across the literature. We highlight some of these reasons.

It is apparent from our analysis that assumptions about the functional form of probability weighting function have important implications. We found that different functional forms yielded remarkably different probability weighting functions. For example, we estimated the more flexible two-parameter functions to be concave, while the one-parameter probability weighting function was essentially linear, suggesting no probability distortions. Thus, selecting the appropriate functional form is important and can dramatically affect the results, especially if probability distortions are not in the traditionally assumed S-shape. Our findings should caution researchers to either use the most flexible probability weighting functions or to estimate a range of them.

Most importantly, we showed that the assumption in prospect theory, — the probability weighting function is static —, is violated at least in our captive macaques. This unambiguous observation was supported by earlier findings, in which the same animals exhibit different probability distortions under different experimental conditions (27, 33), one of which showed the history effect of gamble on both utility and probability distortion (33). Our findings indicate that animal behavior is modelled better when we allow their perceptions of probability to dynamically adjust according to the reward prediction error, a critical algorithmic feature embedded in the nervous system (37, 42). Thus, it is worthy to ask whether our dynamic prospect theory outperform to model gambling behavior across different species including humans.

One of the main challenges of studying probability distortions in humans is the number of choices needed to reliably estimate probability weighting and utility at the individual level. For this reason, experimental tasks often include choices with large-reward low-probability lotteries (12), like a 1% chance of winning $500. Rewards this large are never included in monkey experiments, suggesting that it remains unclear how monkeys behave when faced with small probabilities of really large gains. This may affect the estimated probability distortions since the lower/upper limits largely affect the parameter estimation. However, this experimental design in human studies introduces a major limitation: the negative correlation between the reward magnitudes can affect estimates, opposite to the challeng for reliable parameter estimation. To overcome this issue, we used a dataset that includes a very large number of choices from each individual and payoff matrix fully orthogonalized the reward magnitudes and probabilities. These features allow us to estimate many free parameters reliably but are not feasible for human studies where the duration of sessions and attrition are major concerns in experimental design.

In short, probability distortion is not static rather dynamic, contradicting an assumption in the prospect theory.

### Utility of Monkeys

Most previous studies have found that monkeys have a convex (24, 26, 27) or concave (28, 30-32) utility over rewards in the gain domain. Monkeys in our study, like human decision-makers, were estimated to have a concave utility function. A possible explanation for the convexity is that those studies did not account for the possibility of optimistic probability distortions after lucky wins. If we analyzed our data either assuming no probability distortions or the inflexible one-parameter probability weighting function, each monkey was erroneously defined to have a convex utility or to have a much less concave utility, than when the probability distortions were appropriately accounted for. Due to this limitation, the overweighting probability after lucky wins can be captured through a convex utility function without flexible probability weighting functions. To reliably identify the shape of these functions, the utility function and the probability distortions both need to be estimated, in case the convexity in the utility function differs from the curvature typically estimated in the human decision-makers.

Several aspects distinguish experiments with monkeys from typical studies with humans. Monkeys make many choices for which they obtain immediate rewards that are usually very small (less than 0.3 ml of some liquid), while humans usually make fewer choices for larger rewards. Both of these features are prone to driving monkeys to take more risks than humans in typical experimental conditions. Consistent with this idea, Schultz et al. found that when large rewards were used in a single trial (more than 0.8 ml) the shape of monkeys’ utility function was concave (29). The same phenomenon was observed in our previous experiment (30). In other experimental conditions similar to natural foraging, the utility function of the monkeys was concave (28). Comparing this with humans, Hayden and Platt (59) found that human decision-makers can also display risk-seeking and convex utility functions for rewards in both juice and monetary forms, under conditions resembling those in monkey experiments. If our findings extend to human choosers, they will be of broad economic significance.

## Materials and methods

### Subjects and Surgical Procedures

Two rhesus monkeys were used (Macaca mulatta, SUN, 7.1 kg, male; Macaca fuscata, FU, 6.7 kg, female). All experimental procedures were approved by the Animal Care and Use Committee of the University of Tsukuba (Protocol No H30.336) and performed in compliance with the US Public Health Service’s Guide for the Care and Use of Laboratory Animals. Each animal was implanted with a head-restraint prosthesis. The subjects performed the cued lottery task for 5 days a week. The subjects practiced the cued lottery task for 10 months, after which they became proficient in choosing lottery options.

### Experimental Procedure

Eye movements were measured using a video camera system at 120 Hz. Visual stimuli were generated by a liquid-crystal display at 60 Hz placed 38 cm from the monkey’s face when seated. Animals performed some blocks of choice trials during visually cued lottery task, with each block containing approximately 30 to 60 trials. In the lottery task, green and blue pie charts indicated reward magnitudes from 0.1 to 1.0 mL, in 0.1 mL increments, and reward probabilities from 0.1 to 1.0, in 0.1 increments, respectively. In total, 100 pie-charts were used. Two pie charts were randomly allocated to the left and right options. Data was gathered 4-5 days per week.

### Calibration of the reward supply system

The precise amount of liquid reward was delivered to the monkeys using a solenoid valve. An 18-gauge tube (0.9 mm inner diameter) was attached to the tip of the delivery tube to reduce the variation across trials. The amount of reward in each payoff condition was calibrated by measuring the weight of water with 0.002 g precision (hence, 2μL) on a single trial basis. This calibration method was the same as previously used in (60).

### Mathematical models

We used standard economic models to estimate the utility function and probability weighting function. We used a standard reinforcement learning model to estimate reward prediction error. See SI Material and Methods for details.

## Acknowledgements

The authors wish to express appreciation to Ryo Tajiri, Masafumi Nejime, Vanessa Sihui, Yoshiko Yabana, and Yuki Suwa for their technical assistance. Monkey FU was provided by NBRP “Japanese Monkeys” through the National Bio Resource Project of the MEXT, Japan. This research was supported by JSPS KAKENHI Grant Number JP:15H05374, 19H05007, the Takeda Science Foundation, Council for Addiction Behavior Studies, Narishige Neuroscience Research Foundation, The Ichiro Kanehara Foundation (H.Y.); JSPS KAKENHI Grant Number JP:26710001, MEXT KAKENHI Grant Number JP: 16H06567 (M.M.); and ARC DP190100489 (A.T.).

## Author Contributions

H.Y. designed the research. H.Y. and Y.I. conducted the experiments. T.K. conducted a part of the experiments. H.Y. and A.T. developed analytic tools. A.T and H.Y. analyzed the data. A.T., H.Y., and J.K. evaluated the results. H.Y. and A.T. wrote the manuscript. A.T., Y.I., T.K., J.K., M.M., and H.Y. edited and approved the final manuscript.

## Figures and Legends

**Fig. S1.**
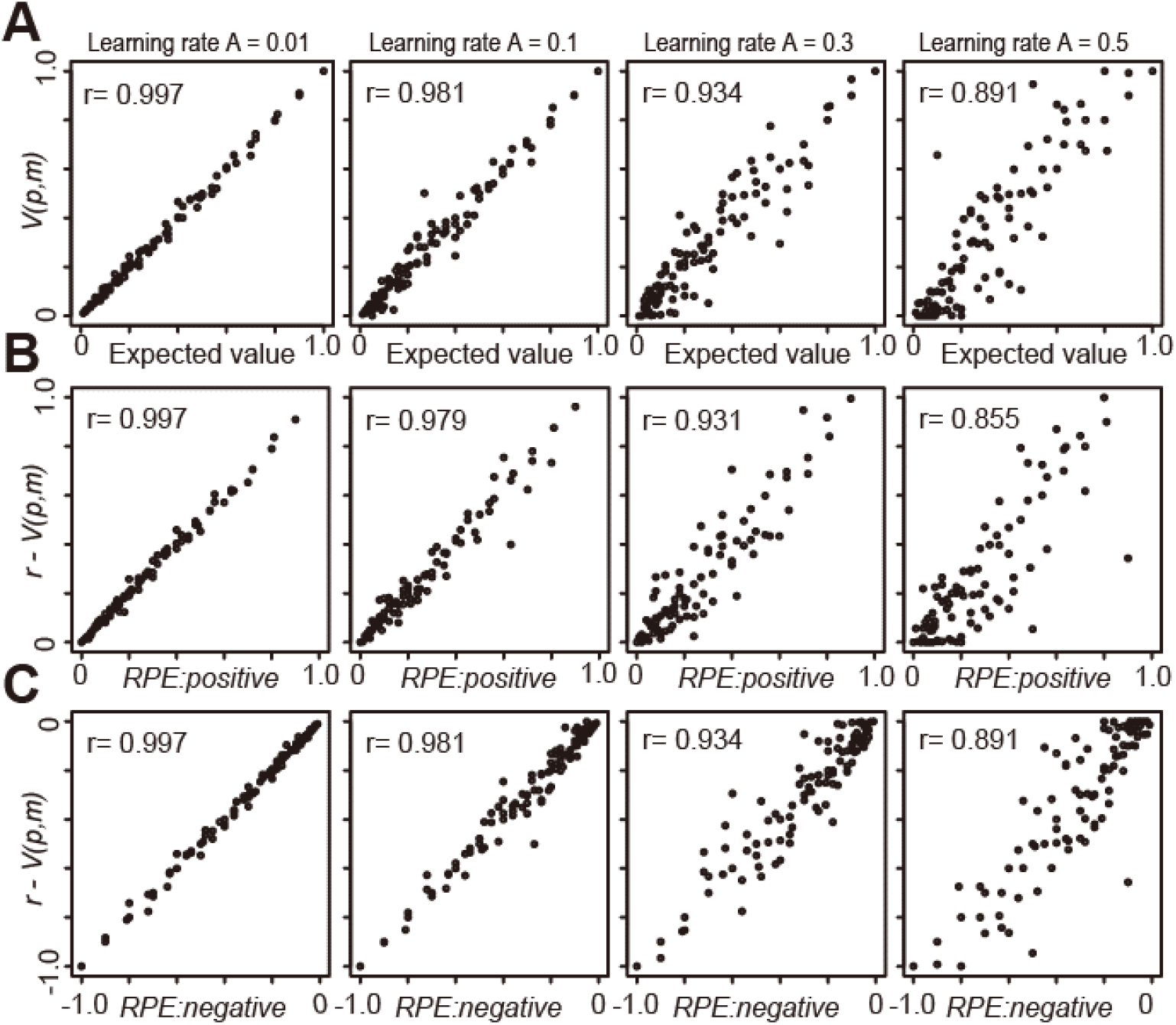
Value function and its RPE estimated by using reinforcement learning model. (A) Plots of the *V(p,m)*_*t=10,000*_ against the expected value defined mathematically, i.e., probability time magnitude. (B) *r - V(p,m)*_*t=10,000*_ after reward against the positive component of RPE, i.e., obtained reward magnitude minus the expected values. (C) *r - V(p,m)*_*t=10,000*_ after no-reward (hence r is zero) against the negative component of RPE, i.e., zero minus the expected values. Plots were made for 100 pie-chart stimuli, as a function of the different learning rate, *A*. Values of r denotes the correlation coefficient.

**Table S1.**
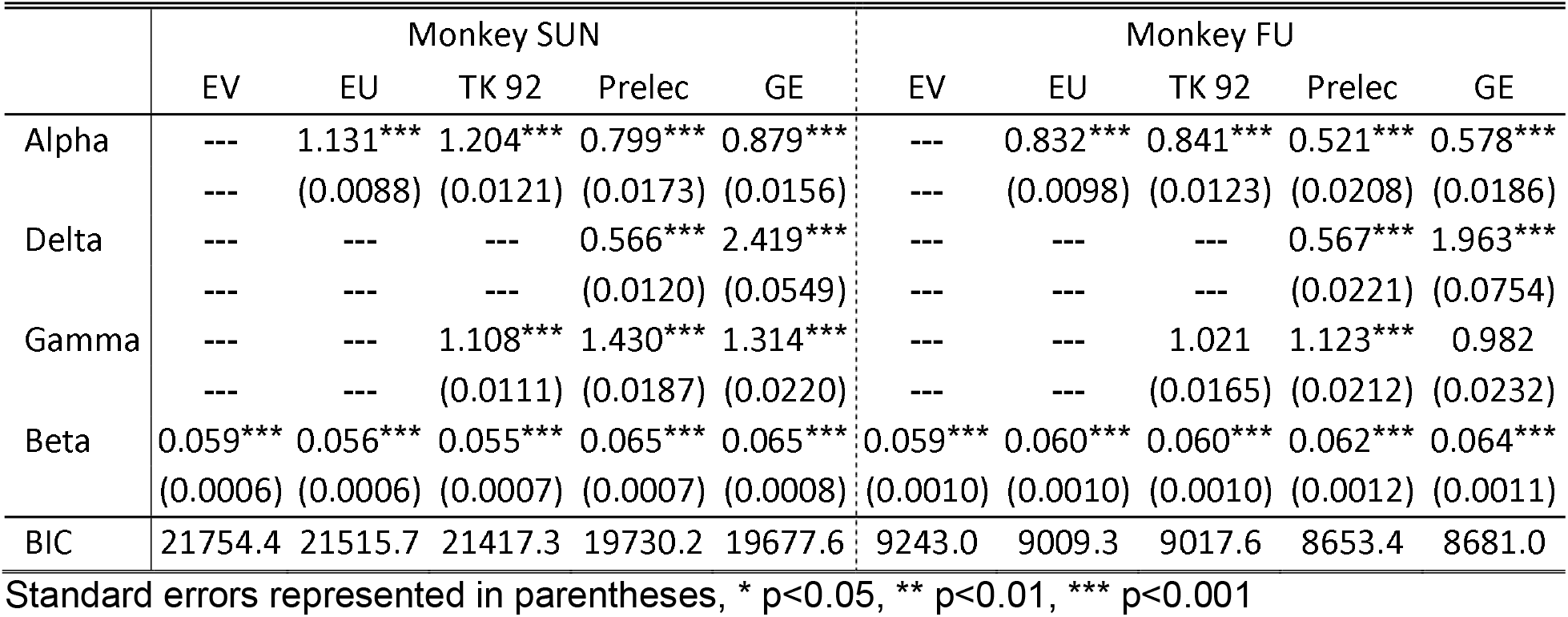
Fit comparison of the standard economic models.

|  | Monkey SUN |  |  |  |  | Monkey FU |  |  |  |  |
| --- | --- | --- | --- | --- | --- | --- | --- | --- | --- | --- |
|  | EV | EU | TK 92 | Prelec | GE | EV | EU | TK 92 | Prelec | GE |
| Alpha | --- | 1.131***<br>(0.0088) | 1.204***<br>(0.0121) | 0.799***<br>(0.0173) | 0.879***<br>(0.0156) | --- | 0.832***<br>(0.0098) | 0.841***<br>(0.0123) | 0.521***<br>(0.0208) | 0.578***<br>(0.0186) |
| Delta | --- | --- | --- | 0.566***<br>(0.0120) | 2.419***<br>(0.0549) | --- | --- | --- | 0.567***<br>(0.0221) | 1.963***<br>(0.0754) |
| Gamma | --- | --- | 1.108***<br>(0.0111) | 1.430***<br>(0.0187) | 1.314***<br>(0.0220) | --- | --- | 1.021<br>(0.0165) | 1.123***<br>(0.0212) | 0.982<br>(0.0232) |
| Beta | 0.059***<br>(0.0006) | 0.056***<br>(0.0006) | 0.055***<br>(0.0007) | 0.065***<br>(0.0007) | 0.065***<br>(0.0008) | 0.059***<br>(0.0010) | 0.060***<br>(0.0010) | 0.060***<br>(0.0010) | 0.062***<br>(0.0012) | 0.064***<br>(0.0011) |
| BIC | 21754.4 | 21515.7 | 21417.3 | 19730.2 | 19677.6 | 9243.0 | 9009.3 | 9017.6 | 8653.4 | 8681.0 |

2 Standard errors represented in parentheses, \* $p < 0.05$ , \*\* $p < 0.01$ , \*\*\* $p < 0.001$

**Table S2.** Additional evidence of concave probability distortions. Results of a logistic regression analysis with an indicator variable for whether the monkey selected target 1 (right) in a trial. m_i_ and p_i_ respectively denote the reward magnitude and probability of receiving reward in target (1: right, 2: left). p_1_<=0.3 (p_1_>=0.7) is an indicator variable equal to one if p_1_ is smaller than or equal to 0.3 (larger than or equal to 0.7).

|  | Monkey SUN | Monkey FU |
| --- | --- | --- |
| $p_1 \leq 0.3$ | -1.656***<br>(0.0681) | -0.886***<br>(0.1017) |
| $p_1 \geq 0.7$ | -1.098***<br>(0.0666) | -0.620***<br>(0.1016) |
| $p_1$ | 6.764***<br>(0.1503) | 7.541***<br>(0.2343) |
| $m_1$ | 7.672***<br>(0.0865) | 6.762***<br>(0.1239) |
| $p_2$ | -6.215***<br>(0.0776) | -7.391***<br>(0.1297) |
| $m_2$ | -7.044***<br>(0.0826) | -6.635***<br>(0.1231) |
| Constant | -0.034<br>(0.0870) | 0.335*<br>(0.1324) |
| N | 44883 | 19292 |

Standard errors represented in parentheses
\* $p < 0.05$ , \*\* $p < 0.01$ , \*\*\* $p < 0.001$

**Table S3.**
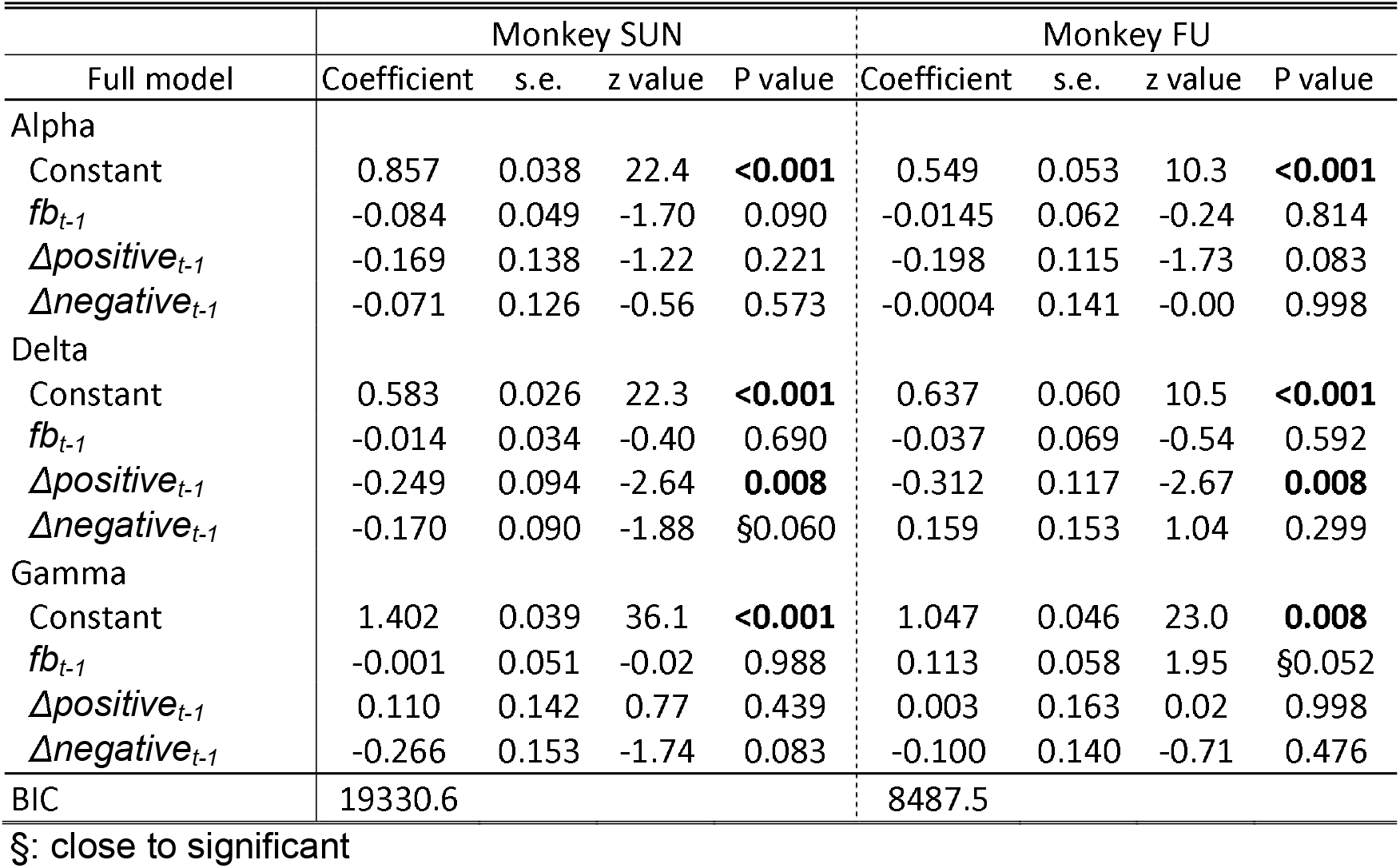
Estimated parameters in full combined model.

**Table S4.**
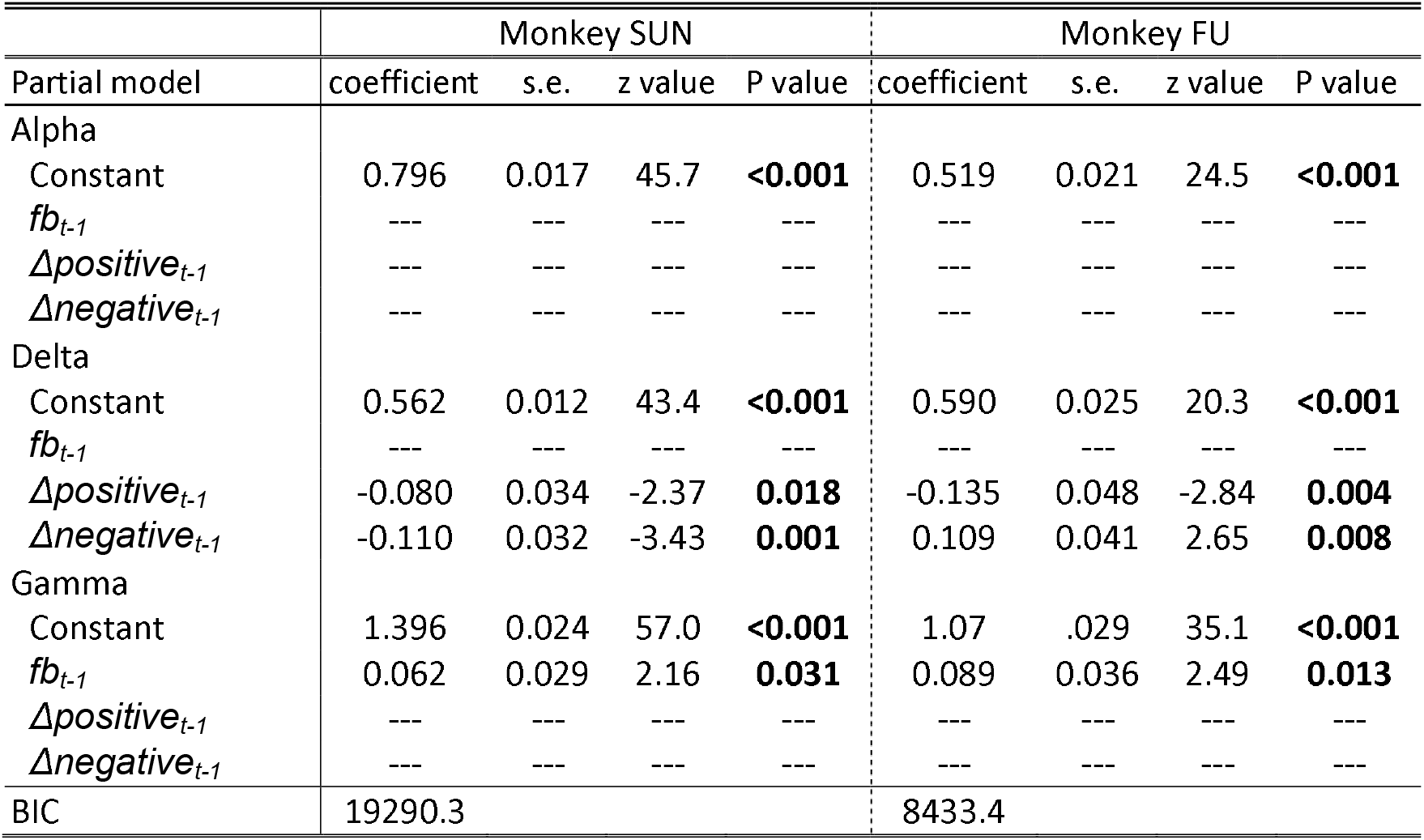
Estimated parameters in the partially combined model based on model selection.

## Supplement: Methods

### Cued lottery tasks

Animals performed one of the two visually cued lottery tasks: a single cue task or a choice task.

### Single cue task

At the beginning of each trial, the monkeys had 2 seconds to align their gaze to within 3° of a 1°-diameter gray central fixation target. After fixating for 1 second, an 8° pie chart providing information about the probability and magnitude of rewards was presented for 2.5 s at the same location as the central fixation target. Magnitude and probability were indicated by numbers of the green and blue pie chart segments, respectively. The pie chart was then removed and 0.2 seconds later, a 1 kHz and 0.1 kHz tone of 0.15 s duration indicated reward and no-reward outcomes, respectively. The high tone preceded reward delivery by 0.2 s. The low tone indicated that no reward was delivered. The animals received a liquid reward as indicated by the number of the green pie chart segments with the probability indicated by the number of the blue pie chart segements. An inter-trial interval of 4 to 6 seconds followed each trial.

### Choice task

At the beginning of each trial, the monkeys had 2 seconds to align their gaze to within 3° of a 1°-diameter gray central fixation target. After fixating for 1 second, two peripheral 8° pie charts providing information about the probability and magnitude of rewards for each of the two target options were presented for 2.5 seconds at 8° to the left and right of the central fixation location. Gray 1° choice targets appeared at these same locations. After a 0.5 second delay, the fixation target disappeared, cueing saccade initiation. The monkeys allowed 2 seconds to make their choice by shifting their gaze to either target within 3° of the choice target. A 1 kHz and 0.1 kHz tone sounded for 0.15 seconds to denote reward and no-reward outcomes respectively. The animals received a liquid reward as indicated by the number of the green pie chart segments of the chosen target with the probability indicated by the number of the blue pie chart segments. An inter-trial interval of 4 to 6 seconds followed each trial.

### Payoff, block structure, and data collection

Green and blue pie charts respectively indicated reward magnitudes from 0.1 to 1.0 mL, in 0.1 mL increments, and reward probabilities from 0.1 to 1.0, in 0.1 increments. A total of 100 pie chart combinations were used. In the single cue task, each pie chart was presented once in a random order, allowing monkeys to experience all 100 lotteries within a certain time period. In the choice task, two pie charts were randomly allocated to the left and right options in each trial. Approximately 30 to 60 trial blocks of the choice task were sometimes interleaved with 100 to 120 trial blocks of the single cue task.

## Statistical Analysis

### Economic models

We estimated parameters of the utility and probability weighting functions within a random utility framework. Specifically, a lottery *L(p,m)* denoted a gamble that pays *m* (magnitude of the offered reward in ml) with probability *p* and 0 otherwise. We assumed a popular constant relative risk attitude (CRRA, also known as power) utility function, *U(m)= m*^*α*^, and considered various previously proposed probability weighting functions. We assumed three subjective probability functions used in the prospect theory, *w(p)*; TK92 (1): *p*^*γ*^ / (*p*^*γ*^+(1-*p*)^*γ*^)^1/*γ*^; Prelec (2): *exp(-δ (-log p)* ^*γ*^); GE (3): *δp*^*γ*^ / (*δp*^*γ*^+(1-*p*)^*γ*^). We assumed that subjective probabilities and utilities are integrated multiplicatively per the standard economic theory, yielding the expected subjective utility function *V(p,m) = w(p) m*^*α*^.

The probability of the monkey choosing the lottery on the right side (*L*_*R*_) instead of the lottery on the left side (*L*_*L*_) was estimated using a logistic choice function:

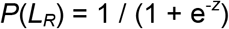

where *Z* = *β* × (*V(L*_*R*_) - *V(L*_*L*_)) and the free parameter *β* indicates the degree of stochasticity observed in choice. We fit the data by maximizing log-likelihood and choose the best structural model to describe the monkeys’ behavior using the Bayesian information criterion (BIC) (4).

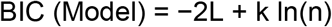

Where, L is the maximum log-likelihood of the model, k is the number of free parameters, and n is the sample size of the model.

In each fitted model, whether *α, γ*, and *δ* were significantly different from zero was determined by a one-sample t-test at P < 0.05. We only used BIC since the dataset was large, meaning that the additional parameter penalty was small if we use AIC.

### Reinforcement learning model

We simulated monkey behavior using a standard temporal difference (TD) learning model (5). Let *V(m,p)*_*t*_ represent the reward prediction from the chosen lottery option in trial *t*. Let *r*_*t*_ be the reward amount received in trial *t* in mL if the trial is rewarded, and 0 if the trial is not rewarded. In a trial *t*, upon receiving the actual reward (or not), the monkeys updated their reward prediction *V(m,p)* _*t+1*_ for future trials using the TD model as follows:

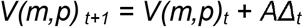

where *A* is the learning rate (0 ≤ *A* ≤ 1), and *Δ*_*t*_ is the reward prediction error. This reward prediction error is defined as:

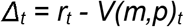

Assuming *V(m,p)*_*0*_ *= 0*, we simulated the TD algorithm for 10,000 experimental trials for each lottery using different values of the learning rate *A*. In Figure S1A, we plot

*V(m,p)*_*10,000*_, which is the valuation arrived at by the algorithm for each lottery against its expected value. Ultimately, the algorithm values the lotteries at close to their expected values, especially for low learning rates. The reward prediction error estimated from the TD algorithm was close to the obtained rewards minus expected values. The *V(m,p)* for the chosen target is updated if the monkeys make chooses that target in trial t.

### Combined models

We calculate the reward prediction error as the difference between the reward received and the expected value of the chosen lottery in the combined model. Therefore, by design, the reward prediction error was positive (*Δpositive*) in our experiment after the receipt of the reward and negative (*Δnegative*) after no reward was given. As shown in the main text, we defined the combined model *V(p,m)*_*t*_ as:

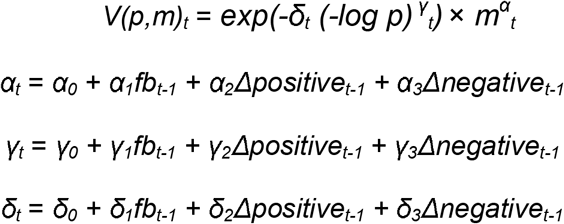

Where *fb*_*t-1*_ is scalar values in the reward (1) and no-reward (0) in the previous trials. *Δpositive*_*t-1*_ is reward amount received minus expected value in the previous reward trials, otherwise zero. *Δnegetive*_*t-1*_ is zero minus expected value in the previous no-reward trials, otherwise zero. In this full combined model, best fitting parameters were estimated based on the BIC value. Whether *α*_*0*_, *γ*_*0*,_ and *δ*_*0*_ were significantly different from 0 was determined by a one-sample t-test using P < 0.05. Whether *α*_*1*_ to *α*_*3*_, *γ*_*1*_ to *γ*_*3*_, *δ*_*1*_ to *δ*_*3*_ were significantly different from one was also determined by one-sample t-test at P < 0.05.

In the partially combined model, we assumed that *V(m,p)*_*t*_ from the full combined model equation is updated by the reward prediction error *Δ*_*t*_ via some combination of those free parameters, *α, γ*, and *δ* as above. We then selected one combination of the free parameters that showed the smallest BIC value among all the combinations of *α*_*0*_ to *α*_*3*_, *γ*_*0*_ to *γ*_*3*_, *δ*_*0*_ to *δ*_*3*_. Note that the full model includes 12 free parameters for *V(p,m)*_*t*_.

## Footnotes

1 We note that we changed the notation from what is usually used in the reinforcement learning literature because we need to distinguish between the parameters of the reinforcement learning model and prospect theory models.

## References

1. P. W. Glimcher, A. Rustichini, Neuroeconomics: the consilience of brain and decision. Science 306, 447–452 (2004).

2. C. Camerer, G. Loewenstein, G. Prelec Neuroeconomics: How Neuroscience Can Inform Economics Journal of Economic Literature 43, 9–64 (2005).

3. P. W. Glimcher, C. F. Camerer, E. Fehr, R. A. Poldrack, Neuroeconomics: Decision Making and the Brain. P. W. Glimcher, C. F. Camerer, E. Fehr, R. A. Poldrack, Eds. (Elsevier, New York, 2008).

4. D. Kahneman, A. Tversky, Prospect theory: An analysis of decisions under risk. Econometrica 47, 313–327 (1979).

5. R. S. Sutton, A. G. Barto, Reinforcement Learning (The MIT press, Cambridge, 1998).

6. B. Garcia, F. Cerrotti, S. Palminteri, The description-experience gap: a challenge for the neuroeconomics of decision-making under uncertainty. Philos Trans R Soc Lond B Biol Sci 376, 20190665 (2021).

7. J. Von Neumann, O. Morgenstern, Theory of Games and Economic Behavior. (Princeton Univ. Press, New Jersey, 1944).

8. P. A. Samuelson, The Problem of Integrability in Utility Theory. Economica 17, 355–385 (1950).

9. H. S. Houthakker, Revealed Preference and the Utility Function. Economica 17, 159–174 (1950).

10. L. J. Savage, The Foundations of Statistics (John Wiley and Sons, New. York, 1954).

11. K. J. Arrow, Essays in the Theory of Risk Bearing. (Markham, Chicago, 1971).

12. G. Wu, R. Gonzalez, Curvature of the Probability Weighting Function. Management Science 42, 1676–1690 (1996).

13. M. Bdellaoui, Parameter-Free Elicitation of Utility and Probability Weighting Functions. Management Science 46, 1497–1512. (2000).

14. W. Harbaugh, K. Krause, L. Vesterlund, Risk attitudes of children and adults: Choices over small and large probability gains and losses. Experimental Economics 5, 53–84. (2002).

15. G. W. Harrison, E. E. Rutstrom, Expected utility theory and prospect theory: One wedding and a decent funeral. Experimental Economics 12, 133–158. (2009).

16. A. Bruhin, H. Fehr-Duda, T. Epper, Risk and Rationality: Uncovering Heterogeneity in Probability Distortion. Econometrica 78, 1375–1412. (2010).

17. H. Fehr-Duda, T. Epper, A. Bruhin, R. Schubert, Risk and rationality: The effects of mood and decision rules on probability weighting. Journal of Economic Behavior and Organization 78, 14–24. (2011).

18. P. N. Tobler, G. I. Christopoulos, J. P. O’Doherty, R. J. Dolan, W. Schultz, Neuronal distortions of reward probability without choice. J Neurosci 28, 11703–11711. (2008).

19. M. Hsu, I. Krajbich, C. Zhao, C. F. Camerer, Neural response to reward anticipation under risk is nonlinear in probabilities. J Neurosci 29, 2231–2237 (2009).

20. D. Stephens, J. Krebs, Foraging Theory (Princeton Univ. Press, New Jersey, 1986).

21. T. Caraco, S. Martindale, T. S. Whitham, An empirical demonstration of risk-sensitive foraging preferences. Animal Behaviour 28, 820–830 (1980).

22. F. Brito e Abreu, A. Kacelnik, Energy budgets and risk-sensitive foraging in starlings. Behavioral Ecology 8, 338–345 (1999).

23. E. U. Weber, S. Shafir, A. R. Blais, Predicting risk sensitivity in humans and lower animals: risk as variance or coefficient of variation. Psychol Rev 111, 430–445 (2004).

24. A. N. McCoy, M. L. Platt, Risk-sensitive neurons in macaque posterior cingulate cortex. Nat Neurosci 8, 1220–1227 (2005).

25. B. Y. Hayden, M. L. Platt, Temporal discounting predicts risk sensitivity in rhesus macaques. Curr Biol 17, 49–53 (2007).

26. W. R. Stauffer, A. Lak, P. Bossaerts, W. Schultz, Economic choices reveal probability distortion in macaque monkeys. J Neurosci 35, 3146–3154 (2015).

27. S. Farashahi, H. Azab, B. Hayden, A. Soltani, On the Flexibility of Basic Risk Attitudes in Monkeys. J Neurosci 38, 4383–4398 (2018).

28. B. R. Eisenreich, B. Y. Hayden, J. Zimmermann, Macaques are risk-averse in a freely moving foraging task. Sci Rep 9, 15091 (2019).

29. W. Genest, W. R. Stauffer, W. Schultz, Utility functions predict variance and skewness risk preferences in monkeys. Proc Natl Acad Sci U S A 113, 8402–8407 (2016).

30. H. Yamada, A. Tymula, K. Louie, P. W. Glimcher, Thirst-dependent risk preferences in monkeys identify a primitive form of wealth. Proc Natl Acad Sci U S A 110, 15788–15793 (2013).

31. S. Ferrari-Toniolo, P. M. Bujold, F. Grabenhorst, R. Baez-Mendoza, W. Schultz, Non-human primates satisfy utility maximization in compliance with the continuity axiom of Expected Utility Theory. J Neurosci 10.1523/JNEUROSCI.0955-20.2020 (2021).

32. A. Nioche, S. Bourgeois-Gironde, T. Boraud, An asymmetry of treatment between lotteries involving gains and losses in rhesus monkeys. Sci Rep 9, 10441 (2019).

33. S. Ferrari-Toniolo, P. M. Bujold, W. Schultz, Probability Distortion Depends on Choice Sequence in Rhesus Monkeys. J Neurosci 39, 2915–2929 (2019).

34. A. Nioche et al., The adaptive value of probability distortion and risk-seeking in macaques’ decision-making. Philos Trans R Soc Lond B Biol Sci 376, 20190668 (2021).

35. P. R. Montague, P. Dayan, T. J. Sejnowski, A framework for mesencephalic dopamine systems based on predictive Hebbian learning. J Neurosci 16, 1936–1947 (1996).

36. W. Schultz, P. Dayan, P. R. Montague, A neural substrate of prediction and reward. Science 275, 1593–1599 (1997).

37. J. C. Houk, J. L. Adams, A. G. Barto, Models of Information Processing in the Basal Ganglia. J. C. Houk, J. L. Davis, D. G. Beiser, Eds. (The MIT Press, Cambridge, 1995), pp. 249–270.

38. J. O’Doherty et al., Dissociable roles of ventral and dorsal striatum in instrumental conditioning. Science 304, 452–454 (2004).

39. D. J. Barraclough, M. L. Conroy, D. Lee, Prefrontal cortex and decision making in a mixed-strategy game. Nat Neurosci 7, 404–410 (2004).

40. N. D. Daw, Y. Niv, P. Dayan, Uncertainty-based competition between prefrontal and dorsolateral striatal systems for behavioral control. Nat Neurosci 8, 1704–1711 (2005).

41. K. Samejima, Y. Ueda, K. Doya, M. Kimura, Representation of action-specific reward values in the striatum. Science 310, 1337–1340 (2005).

42. A. S. Lowet, Q. Zheng, S. Matias, J. Drugowitsch, N. Uchida, Distributional Reinforcement Learning in the Brain. Trends Neurosci 43, 980–997 (2020).

43. K. Enomoto, N. Matsumoto, H. Inokawa, M. Kimura, H. Yamada, Topographic distinction in long-term value signals between presumed dopamine neurons and presumed striatal projection neurons in behaving monkeys. Sci Rep 10, 8912 (2020).

44. H. Yamada, Y. Imaizumi, M. Matsumoto, Neural Population Dynamics Underlying Expected Value Computation. J Neurosci 41, 1684–1698 (2021).

45. J. D. Hey, C. Orme, Investigating Generalizations of Expected Utility Theory Using Experimental Data. Econometrica 62, 1291–1326 (1994).

46. Y. Wang et al., Neural substrates of updating the prediction through prediction error during decision making. Neuroimage 157, 1–12 (2017).

47. D. Prelec, The Probability Weighting Function. Econometrica 66, 497–527 (1998).

48. W. M. Goldstein, H. J. Einhorn, Expression theory and the preference reversal phenomena. Psychological Review 94, 236–254 (1987).

49. H. Seo, X. Cai, C. H. Donahue, D. Lee, Neural correlates of strategic reasoning during competitive games. Science 346, 340–343 (2014).

50. M. Matsumoto, K. Matsumoto, H. Abe, K. Tanaka, Medial prefrontal cell activity signaling prediction errors of action values. Nat Neurosci 10, 647–656 (2007).

51. H. Inokawa, N. Matsumoto, M. Kimura, H. Yamada, Tonically Active Neurons in the Monkey Dorsal Striatum Signal Outcome Feedback during Trial-and-error Search Behavior. Neuroscience 446, 271–284 (2020).

52. H. Zhang, X. Ren, L. T. Maloney, The bounded rationality of probability distortion. Proc Natl Acad Sci U S A 117, 22024–22034 (2020).

53. A. Tymula, P. Glimcher, Expected Subjective Value Theory (ESVT): A Representation of Decision Under Risk and Certainty. SSRN (2020).

54. H. Yamada, Hunger enhances consistent economic choices in non-human primates. Sci Rep 7, 2394 (2017).

55. L. Pompilio, A. Kacelnik, S. T. Behmer, State-dependent learned valuation drives choice in an invertebrate. Science 311, 1613–1615 (2006).

56. A. Fujimoto, T. Minamimoto, Trait and State-Dependent Risk Attitude of Monkeys Measured in a Single-Option Response Task. Front Neurosci 13, 816 (2019).

57. M. Abdellaoui, Parameter-Free Elicitation of Utility and Probability Weighting Functions.. Management Science 46, 1497–1512. (2000).

58. A. Bruhin, H. Fehr-Duda, T. Epper, Risk and Rationality: Uncovering Hetero-geneity in Probability Distortion. Econometrica 78, 1375–1412. (2010).

59. B. Y. Hayden, M. L. Platt, Gambling for Gatorade: risk-sensitive decision making for fluid rewards in humans. Anim Cogn 12, 201–207 (2009).

60. H. Yamada, K. Louie, A. Tymula, P. W. Glimcher, Free choice shapes normalized value signals in medial orbitofrontal cortex. Nat Commun 9, 162 (2018).

## Supplementary references

1. D. Kahneman, A. Tversky, Prospect theory: An analysis of decisions under risk. Econometrica 47, 313–327 (1979).

2. D. Prelec, The Probability Weighting Function. Econometrica 66, 497-527 (1998).

3. W. M. Goldstein, H. J. Einhorn, Expression theory and the preference reversal phenomena. Psychological Review 94, 236–254 (1987).

4. K. Burnham, D. Anderson, Multimodel inference: understanding AIC and BIC in model selection. Sociol. Method Res. 33, 261–304 (2004).

5. R. S. Sutton, A. G. Barto, Reinforcement Learning (The MIT press, Cambridge, 1998).

